# An Integrated Multimodal Model of Alcohol Use Disorder Generated by Data-Driven Causal Discovery Analysis

**DOI:** 10.1101/2020.09.21.306761

**Authors:** Eric Rawls, Erich Kummerfeld, Anna Zilverstand

## Abstract

**Objective:** Alcohol use disorder (AUD) has high prevalence and adverse societal impacts, but our understanding of the factors driving AUD is hampered by a lack of studies that describe the complex multifactorial mechanisms driving AUD.

**Methods:** We used Causal Discovery Analysis (CDA) with data from the Human Connectome Project (HCP; n = 926 [54% female], 22% AUD [37% female]). Our outcome variable was number of AUD symptoms. We applied exploratory factor analysis (EFA) to parse phenotypic measures into underlying constructs, and assessed functional connectivity within 12 resting-state brain networks as an indicator of brain function. We then employed data-driven CDA to generate an integrated model relating phenotypic factors, fMRI network connectivity, and AUD symptom severity.

**Results:** EFA extracted 18 factors representing the wide HCP phenotypic space (100 measures). CDA produced an integrated multimodal model, highlighting a limited set of causes of AUD. The model proposed a hierarchy with causal influence propagating from brain function to cognition (fluid/crystalized cognition, language & working memory) to social (agreeableness/social support) to affective/psychiatric function (negative affect, low conscientiousness/attention, externalizing symptoms) and ultimately AUD severity. Every edge in the model was present at *p* < .001, and the SEM model overall provided a good fit (RMSEA = .06, Tucker-Lewis Index = .91).

**Conclusions:** Our data-driven model confirmed hypothesized influences of cognitive and affective factors on AUD, while underscoring that traditional addiction models need to be expanded to highlight the importance of social factors, amongst others. Results further demonstrated that it is possible to extract a limited set of causal factors of AUD, which can inform future research aimed at tracking factors that dynamically predict alcohol use trajectories. Lastly, the presented model identified potential treatment targets for AUD, including neuromodulation of the frontoparietal network, cognitive/affective interventions, and social interventions.

## 1 Introduction

Lifetime incidence of alcohol use disorder (AUD) is as high as 29-30% (1,2), with alcohol use being a leading cause of death [3 million in the United States, (3); 5.3% of all deaths worldwide; (4)]. Success rates for quitting drinking in AUD are low [30-40%; (5,6)], which has been attributed to the multi-causality of the mechanisms underlying AUD and the need for more targeted treatments (7). However, the development of targeted interventions for AUD is hampered by a lack of studies investigating multifactorial mechanisms driving AUD.

Early theories of addiction maintenance proposed single key mechanisms, such as allostasis (8), or hedonic signaling (9). These early theories have given way to multifactorial models of addiction, such as the “three-stage cycle” model (10), which proposes that negative affect, incentive salience, and executive function are functional domains involved in addiction. There is a great deal of empirical support for the involvement of these three domains. The three domains have been mapped onto corresponding personality profiles that confer addiction risk (11) and have been used to develop a set of proposed neuroclinical assessment tools (12,13) that were successfully applied to AUD (14). However, a three-domain model is far from encompassing the entire phenotypic space that contributes to AUD. The NIMH RDoC (15,16) proposed 23 functional domains underlying psychopathology, recognizing a need for multivariate models that incorporate increasing knowledge of the many functional domains contributing to psychiatric dysfunction. A recent consensus paper on a multivariate assessment approach for addiction, identified another seven “addiction-specific” domains in addition to the RDoC domains (17). Critically, in all of these approaches, the proposed functional domains were identified by expert consensus and therefore might not exactly match the true underlying domains that exist in the data. For example, in an exploratory analysis of a large public dataset, Van Dam et al. (18) derived seven phenotypic factors that only partially mapped onto RDoC domains, but predicted psychiatric distress. A more recent addiction theory (19) identified ten domains contributing to maladaptive decision-making in addiction. A systematic review of neuroimaging studies in addicted populations, implicated the involvement of at least six different neurobiological mechanisms in AUD (20). These recent developments underscore the need for data-driven, multivariate analysis methods capable of fully examining and describing the large phenotypic space underlying addiction, if we are to understand the central question of how multi-causal factors underlie the maintenance and escalation of alcohol use.

In the current study, we leveraged the deep behavioral and psychiatric phenotyping (21) and high-resolution neuroimaging data (22) from the Human Connectome Project (HCP) (23). Using data from nearly 1000 participants we first derived a set of data-driven domains underlying the full range of phenotypic functioning measured in the HCP dataset. We extracted whole-brain connectivity metrics from 12 data-derived resting-state fMRI networks (24) to measure individual neurobiological differences. To examine the relationships between fMRI network connectivity, phenotypic domains, and AUD symptom severity, we applied Causal Discovery Analysis (CDA), a class of machine learning techniques that learns causal models from input data. These methods search the enormous set of possible structural models and return a graph representing estimated causal relationships in the data. The particular method we applied, Greedy Fast Causal Inference (25), uses conditional dependence relations to discover when unmeasured variables confound the relationships between measured variables, making this method particularly powerful for real-world data sets that cannot possibly capture every variable of interest. By 1) deriving data-driven domains encompassing the whole phenotypic space measured in HCP, 2) extracting whole-brain network connectivity profiles, and 3) applying CDA to the resulting phenotypic and neurobiological domains, we generated an integrated, multimodal causal model of neurobehavioral factors contributing to AUD symptom severity.

## 2 Methods

### 2.1 Subjects

We analyzed data from the final release of the WU-Minn Human Connectome Project (n = 1206, aged 22 – 35, 54% female). All subjects provided written informed consent at Washington University. The causal discovery analysis included all subjects who had complete data from all modalities (phenotypic n = 933, resting-state fMRI n = 1085, final n = 926). Subjects with AUD comprised 22% of the included sample, and 37% of subjects with AUD were female. See Table 1 for demographic characteristics of the included sample.

**Table 1.** Demographic characteristics of the final sample (n = 926).

| Demographics | Options | Total N | AUD | Control | AUD - Control Difference |
| --- | --- | --- | --- | --- | --- |
| **Gender** | M | 428 | 128 | 300 | $\chi^2 = 28.74, p < .001$ |
|  | F | 498 | 76 | 498 |  |
| **Race** | White | 700 | 166 | 534 | $\chi^2 = 19.56, p = .002$ |
|  | Black/African-American | 130 | 15 | 115 |  |
|  | Asian/Nat. Hawaiian/Other Pacific Islander | 57 | 8 | 49 |  |
|  | Other | 39 | 15 | 24 |  |
| **Age** | Mean | 28.84 | 28.65 | 28.88 | $t(924) = 0.80, p = .43$ |
|  | Standard Deviation | 3.69 | 3.38 | 3.74 |  |
| **AUD Symptoms** | 0 | 538 |  |  |  |
|  | 1 | 184 |  |  |  |
|  | 2 | 98 |  |  |  |
|  | 3 | 46 |  |  |  |
|  | 4 | 43 |  |  |  |
|  | 5+ | 17 |  |  |  |
| **AUD Diagnosis** | Yes | 204 |  |  |  |
|  | No | 722 |  |  |  |

### 2.2 Outcome Measure: AUD Symptom Severity

Subjects were assessed for symptoms of alcohol abuse and dependence using the Semi-Structured Assessment for the Genetics of Alcoholism (SSAGA). Symptom count data were provided for DSM-IV-TR alcohol abuse and alcohol dependence. Symptom counts for alcohol abuse were provided as 0, 1, or 2+, and symptom counts for alcohol dependence were provided as 0, 1, 2, or 3+ (i.e. truncated symptom counts were provided). Given the low number of subjects with the highest symptom counts, it is unlikely that including a more fine-grained symptom count would have provided much additional information. It is therefore also likely that participants mostly had mild/moderate AUD severity, and that as the sample was young adults (age 22-35), this sample likely represents an early stage of AUD.

DSM-5 re-categorized alcohol abuse and alcohol dependence into a single disorder (AUD) using the criteria of both alcohol abuse and dependence, with one symptom of alcohol abuse removed (legal problems) and one symptom added (craving). We reconstructed this change by adding alcohol abuse and dependence symptom counts for each subject. Given recent interest in dimensional rather than categorical psychiatric dysfunction, we used the AUD symptom count (severity) as our primary outcome variable.

### 2.3 Behavioral and Self-Report Measures

The HCP dataset contains a wide array of self-report, diagnostic and behavioral measures assessing domains of cognition, emotion, social function, psychiatric dysfunction, and personality. We selected all available measures from these domains for further analysis (100 in total). We included all provided measures, but when provided, we used summary scores rather than item-level or minor scores as long as the summary score encompassed the task construct of interest (26). For example, for the Short Continuous Penn Test, we included the summary statistics of sensitivity and specificity but not more specific scores, and for delay discounting we included the area under the curve for $200 and $40k but not the individual discounting levels. For in-scanner tasks, we included a separate measure for each of the major conditions. If both age/gender-adjusted and unadjusted scores were provided, we included only the adjusted scores. We also excluded all items that were linear combinations of other data. For a list of included variables and excluded variables and descriptive statistics of the included variables see the Supplement.

### 2.4 Factor Analysis of Phenotypic Data

To reduce the phenotypic space measured in the Human Connectome Project to a set of underlying domains, we conducted an exploratory factor analysis (maximum likelihood factor extraction, oblimin rotation to allow for correlated factors) in the entire HCP sample that had complete phenotypic data (n = 933, 53.5% female). EFA was calculated in R using the ‘psych’ package (27). The choice of EFA over similar data reduction schemes such as PCA was made because EFA explicitly accounts for error due to unreliability in measurement (28) unlike PCA (29). Oblimin rotation allows for correlated factors, which is critical to data reduction over a large phenotypic variable space as we expect many factors to be closely related but separable (for example, negative affect and internalizing psychopathology). We used Monte Carlo permutation analysis (parallel analysis) (30) to determine how many factors were statistically significant at *p* < .05 (31). Monte Carlo simulation was also calculated using the ‘psych’ package for R (27).

### 2.5 Resting-State fMRI Acquisition and Preprocessing

High-resolution structural and functional MRI data were collected on a Siemens 3T Connectome Skyra scanner with a 32-channel head coil at Washington University. See Uğurbil et al. (22) for a full description of the acquisition parameters for rfMRI in the HCP database. Structural data were preprocessed as described in Glasser et al. (32), using the most recent version of the HCP preprocessing pipeline (4.1). Briefly, anatomical image preprocessing consisted of bias field and gradient distortion correction, coregistration of T1w and T2w images, and linear and nonlinear registration to MNI space. Cortical surfaces were constructed using FreeSurfer. Surface files were transformed to MNI space, registered to the individual’s native-mesh surfaces, and downsampled.

Functional MRI preprocessing is fully described in Glasser et al. (32). Briefly, volumetric fMRI were subjected to gradient distortion correction, motion correction, and referencing to T1w. All transforms were concatenated and run in a single nonlinear resampling to MNI space. Data were then masked by the PostFreeSurfer brain mask and normalized. This volumetric timeseries was then mapped to a combined cortical surface and subcortical voxel space (“grayordinates”) and smoothed with a 2mm FWHM Gaussian.

Finally, fMRI data were high-pass filtered (FWHM = 2355 s) and cleaned of artifacts using ICA-FIX (33,34). Artifact components and 24 motion regressors (35) were regressed out of the data in a single step. This produced the final ICA-FIX denoised version of the data (https://www.humanconnectome.org/study/hcp-young-adult/document/1200-subjects-data-release) that we used for all analyses.

### 2.6 rfMRI Parcellation and Network Assignment

We parcellated the whole brain, including cortex, subcortex, and cerebellum, into 718 parcels using the Cole-Antecevic parcellation (24), a parcellation scheme that builds on the Glasser et al. (36) multimodal cortex parcellation (360 parcels). We chose the Cole-Antecevic parcellation because while the Glasser parcellation used multiple measures including myelination, rfMRI activity, and anatomical landmarks to delineate a fine-grained map of cortical space, it did not include any subcortical voxels, and did not explicitly assign parcels to large-scale networks using principled statistical methods. The Cole-Antecevic parcellation built on the Glasser parcellation by 1) assigning subcortical and cerebellar voxels to parcels, and 2) by using Louvain community detection to delineate 12 large-scale networks consisting of cortical, subcortical, and cerebellar regions.

### 2.7 Calculation of rfMRI Network Connectivity

For each of the 12 data-derived networks, we computed pairwise Pearson correlations between each pair of parcels in the network. Pearson correlations were transformed to approximate a normal distribution using Fisher’s *z*-transform. Within each of the 12 networks, we then took the average of the parcel-to-parcel correlations to obtain a summary statistic for within-network connectivity (37–41). This procedure therefore summarizes how tightly connected (coherent) the regions comprising each of the 12 networks are with each other. This resulted in 12 average network-level connectivity values for each subject. Figure 1 contains a graphical depiction of whole-brain network assignments.

**Figure 1.**
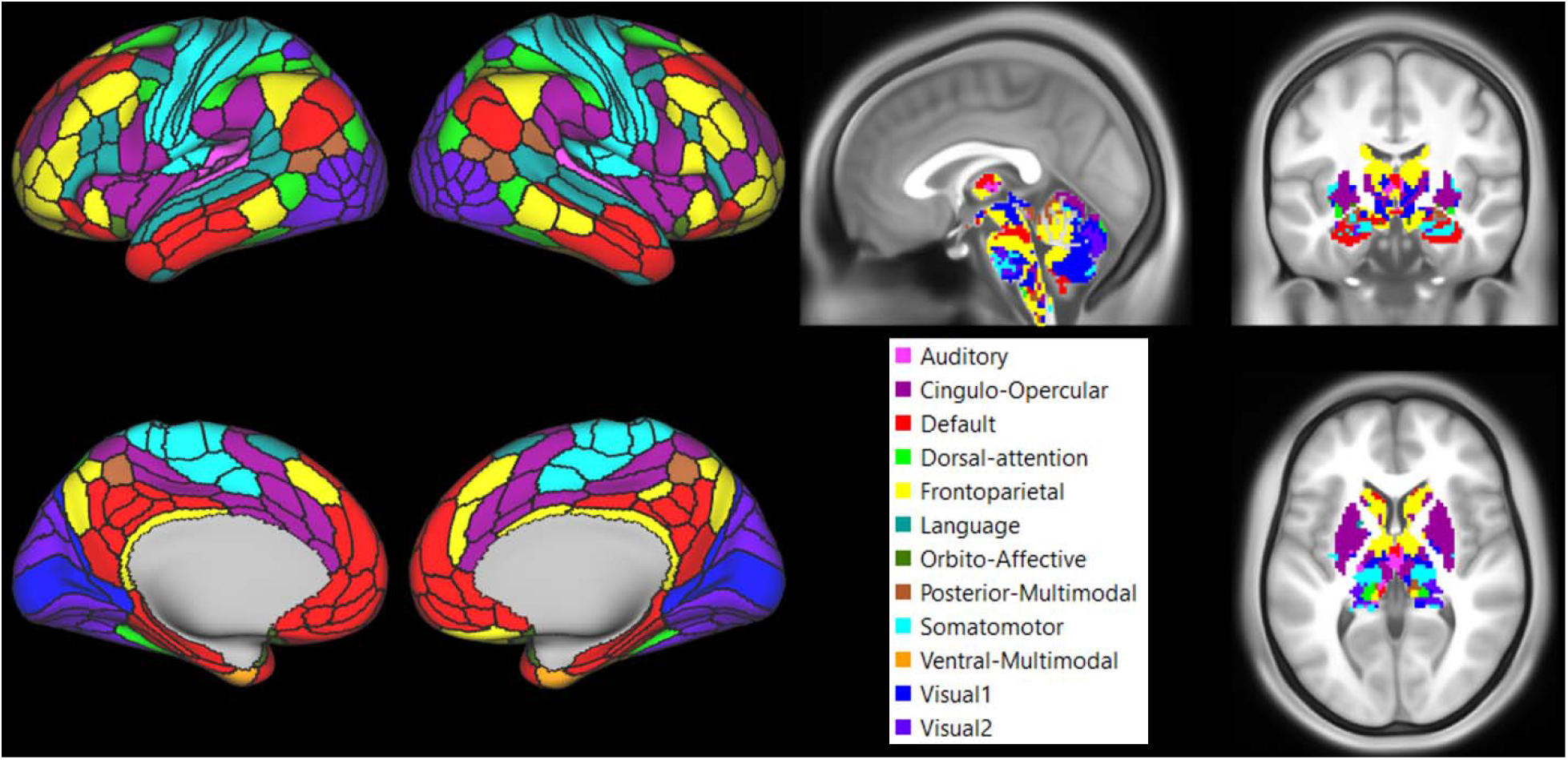
We conducted a whole-brain parcellation and assigned brain parcels to 12 large-scale networks according to the Cole-Antecevic Parcellation (24). This parcellation built on the Glasser multimodal cortical parcellation by including subcortical and cerebellar parcels, and assigned each of the 718 parcels to a large-scale brain network using Louvain community detection.

### 2.8 Causal Discovery Analysis: Greedy Fast Causal Inference (GFCI)

Causal models represent, often graphically, the set of cause-and-effect relationships that are present within a set of data (42). As the number of variables in a dataset increases, so too does the space of possible causal models that could give rise to the observed data, making the problem of identifying which of the potential causal models best fits the observed data very difficult. Causal Discovery Analysis (CDA) uses machine learning to determine which causal models are best supported by the data (43). There are many CDA algorithms that make a wide variety of assumptions and have varying performance characteristics; for review, see Glymour et al. (44).

In the current study, we applied Greedy Fast Causal Inference (GFCI) (25), an accurate and fast algorithm for establishing causal relationships from data even in the presence of unmeasured confounds. GFCI operates in two phases. GFCI begins by searching the space of possible graphs to create a preliminary graph that minimizes a penalized likelihood score, in this case the Bayesian Information Criteria (BIC) (45). This initial search phase is done using the Fast Greedy Equivalence Search method (46). After the initial search phase, the algorithm refines the discovered graph by conducting a series of conditional independence tests. This phase rules out any edges that imply conditional dependencies not borne out by the data (for an example, see *Figure 2*). The most important distinction of GFCI compared to other causal discovery methods is that GFCI can detect confounding factors, and as such is particularly suited to analysis of real-world data, where there is no guarantee that every relevant variable has been measured. The output of the GFCI algorithm is a partial ancestral graph with edge types indicating causal relationships, uncertain relationships, and the presence of unmeasured confounding variables.

**Figure 2.**
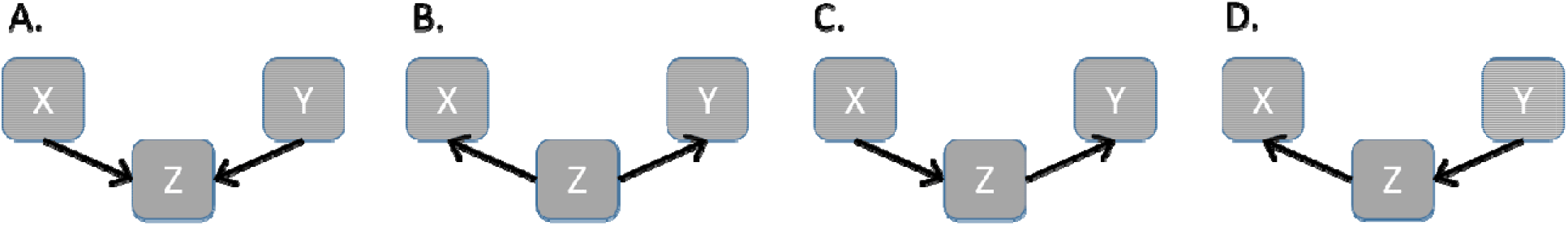
Four different ways that three variables X, Y, and Z could be causally related. Panel **A** is a structure known as a “collider,” in which X and Y both cause Z, but X and Y are not related. In this structure, X and Z are dependent, and Y and Z are dependent, while X and Y are independent. However, when Z is conditioned on (controlled for), X and Y are dependent. Meanwhile, in panel **B**, Z causes both X and Y. In this structure, X and Y are dependent because of their common cause, and are independent when Z is conditioned on. In Panels **C & D**, X and Y are dependent because one causes Z, which then causes the other. In both of these panels, X and Y are independent when Z is conditioned on, as Z is the only link from X to Y. GFCI utilizes conditional independence tests to determine causal direction in graph edges, specifically by identifying “collider” cases in the graph (since these cases imply different conditional dependencies than the other three cases).

GFCI analysis was implemented using Tetrad v6.7, running in Java. Analysis was run with default parameters; that is, using alpha = .01, maximum degree of the graph = 100, and a penalty discount of two. Penalized likelihoods for models were calculated using the Bayesian Information Criteria (45), which is the default model fit index in Tetrad and the most commonly used model fit index in CDA. To recover effect sizes of the edges in the model, we fit a structural equation model (SEM) to the graph structure using the ‘lavaan’ package for R (47). We present the graph GFCI learned from the full dataset with SEM effect sizes for each edge. As an additional analysis of model stability in smaller samples, we also conducted a stability analysis by resampling 90% of the sample (48) without replacement with 2000 repetitions (jackknifing); the resampling stability of each edge is presented in the Supplement.

## 3 Results

### 3.1 Exploratory Factor Analysis: Decomposing the Phenotypic Space of the HCP

Based on the results of Monte Carlo simulation we extracted 18 factors (*p* < .05) from the 100-variable space, which collectively accounted for 47% of common variance. Factor loadings for each item are described in the Supplement. Results of the Monte Carlo permutation test for eigenvalue significance, and percent variance explained per factor, eigenvalues, and cumulative variance are presented in the Supplement. EFA model fit indices indicated good factor separation (RMSEA = .03, Tucker-Lewis Index = .86).

Factors, in order of common variance accounted for, were associated with: 1) Somaticism (high DSM/ASR somaticism, high DSM depression, low PSQI sleep quality), 2) Fluid Cognition (high Raven’s progressive matrices performance), 3) Internalizing (high DSM/ASR anxiety, high DSM depression, high NEO-FFI neuroticism), 4) Gambling Task Reaction Time (slow gambling task reaction time), 5) Conscientiousness/Attention (low DSM ADHD, low ASR attention problems, and high NEO-FFI conscientiousness), 6) Visuospatial Processing (high Penn short line orientation task performance), 7) Social Support (high NIH toolbox friendship, low loneliness, low perceived rejection and perceived hostility, high emotional and instrumental support), 8) Processing Speed (high NIH toolbox flanker total score, fast fMRI emotion task RT), 9) Externalizing (high ASR aggression and rule-breaking, high DSM antisocial, high NIH toolbox aggression), 10) Social Withdrawal (high ASR withdrawal, high DSM avoidance, low NEO-FFI extraversion), 11) Language Task Performance (high fMRI language task story average difficulty, and high math problem accuracy), 12) Relational Task Reaction Time (slow fMRI relational task RT), 13) Delay Discounting (high delay discounting AUC for $200 and $40k), 14) Memory Performance (fMRI N-Back task fast RT and high accuracy, fast Penn word memory RT), 15) Negative Affect (high NIH toolbox anger, fear, sadness and stress), 16) Crystalized IQ (high NIH toolbox English reading and picture vocabulary, high education, and high NEO-FFI openness), 17) Positive Affect (high NIH toolbox life satisfaction, positive affect, and meaning and purpose, and NEO-FFI extraversion), and 18) Agreeableness (low aggression and high NEO-FFI agreeableness).

### 3.2 Causal Discovery of the Neurobehavioral Underpinnings of Alcohol Use Disorder

The output of GFCI is presented in *Figure 3*. The SEM fit to this model indicated a good fit, RMSEA = .06, Tucker-Lewis Index = .91. Recovered edge weights from SEM were presented overlaid on the GFCI graph. Jacknife testing demonstrated the stability of the model (2000 repetitions; see Supplement).

**Figure 3.**
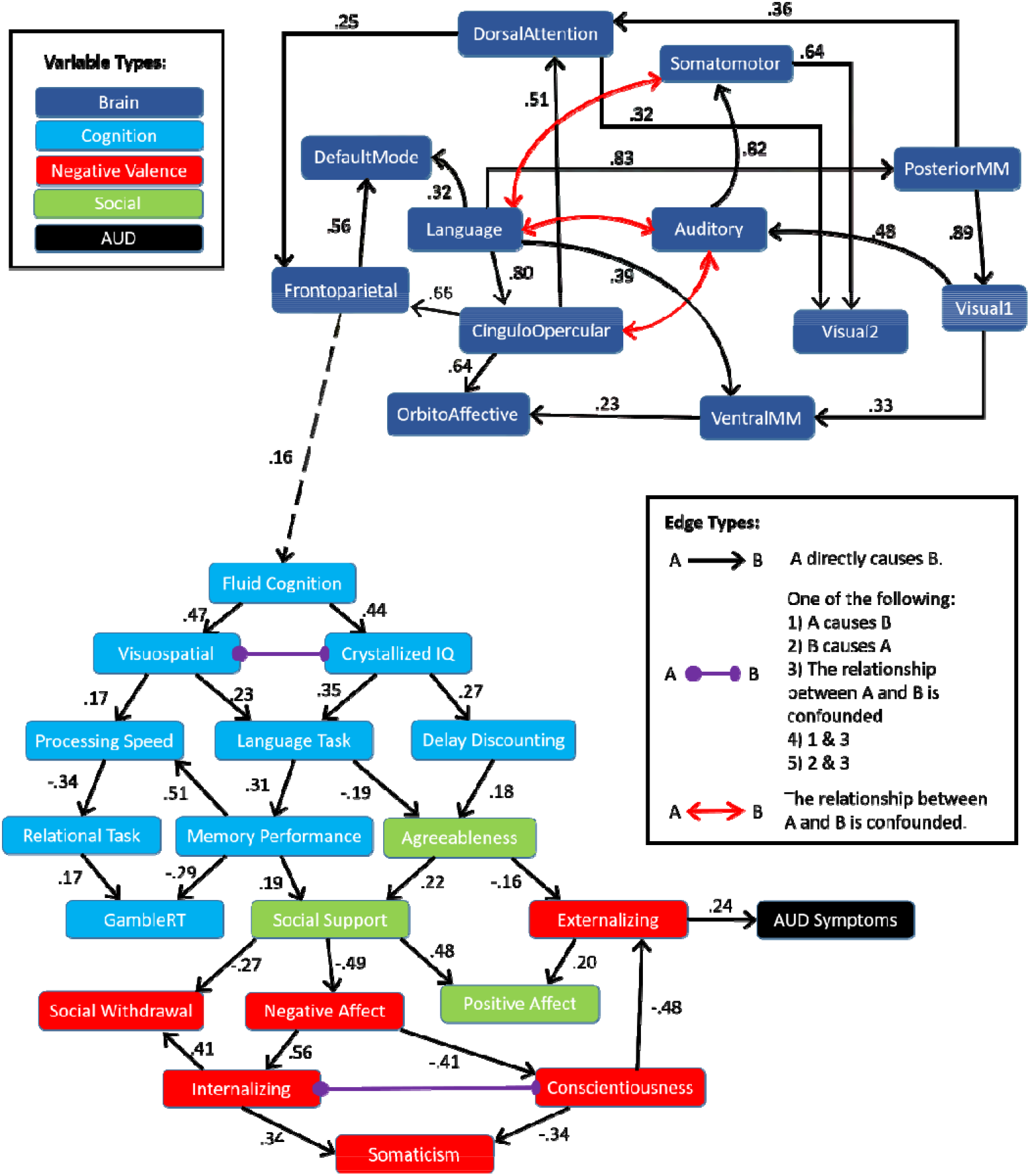
Causal Discovery Analysis of the neurobehavioral determinants of AUD symptom severity in the HCP dataset was done using Greedy Fast Causal Inference (GFCI). GFCI returns a partial ancestral graph (PAG) depicting causal relationships between a set of variables, while assessing for unmeasured third variables in relationships (confounders). Standardized edge weights recovered via SEM are displayed in text next to each edge in the graph.

First, we found that brain network connectivity measures and phenotypic factors largely separated into two interconnected separate clusters. The brain network subgraph indicated several salient points. We found that high connectivity within the language network during rest caused high connectivity within default mode (DMN), cingulo-opercular, and multi-modal sensory association networks - networks that play a central role in self-reflective brain processes. We also found effects indicating causal influences of cingulo-opercular connectivity on attentional processing (cingulo-opercular -> dorsal attention). Finally, we found converging causal influences onto the frontoparietal network (cingulo-opercular/posterior multimodal -> frontoparietal, dorsal attention -> frontoparietal).

Brain connectivity intersected with behavioral phenotypic variables in a link between high frontoparietal connectivity and high Fluid Cognition. From this point, causal influences propagated from Fluid Cognition to Visuospatial Processing and Crystalized IQ, replicating a well-studied effect that individuals high in fluid cognitive ability will also be high in crystalized intelligence (49,50). From there causal influences proceeded to more specific cognitive measures, including Working Memory, Language Task Performance and Delay Discounting. We found a direct link between Crystalized IQ and Delay Discounting, such that individuals higher in Crystalized IQ also exhibited lower (less impulsive) discounting rates. These cognitive measures were then in turn causally linked to affective, social and psychiatric factors. High Language Task performance and less impulsive Delay Discounting caused Agreeableness. Low Working Memory performance and low Agreeableness caused lowered Social Support, and decreased Social Support contributed to increased Negative Affect, increased Social Withdrawal, and decreased Positive Affect. High Negative Affect in turn contributed to higher Internalizing symptoms, and lower Conscientiousness/Attention. Low Conscientiousness/Attention and low Agreeableness caused high Externalizing psychopathology, while high Externalizing psychopathology directly caused increased AUD symptom severity.

Previous hypotheses have particularly focused on the influences of negative affect, incentive salience, and executive function in AUD. Our results support a causal role for cognitive and affective influences on AUD, while expanding our understanding of the complex multifactorial space contributing to AUD.

## 4 Discussion

Early addiction models posited that addiction was due to single key mechanisms (8,9), but modern addiction models have begun to emphasize the multifactorial mechanisms underlying addiction (10,12,17,19,20). In this study, we used a data-driven approach to characterize phenotypic domains in a large community sample, and examined whole-brain network connectivity at a large scale using a data-driven network analysis and parcellation (24). We then modeled large-scale brain and behavioral influences on alcohol use disorder (AUD) symptom severity using causal discovery analysis. Our results shed light on the relationship between brain network connectivity and phenotypic domains in general, as well as providing specific information on how brain and behavioral factors contribute to the severity of AUD, and which could be targeted in treatment.

### 4.1 Expanding the Multifactorial Space of Current Addiction Models

Our factor analysis uncovered a variety of factors that map relatively well onto domains elaborated in RDoC (*Table 2*). For example, we found factors that mapped well onto aspects of the RDoC Cognitive Systems domain, the RDoC Negative Valence Systems domain, and the RDoC Social domain. To assist in interpreting the large-scale domains that our data-driven factors mapped onto, we grouped factors based on their correlations. Interestingly, we found that the Conscientiousness/Attention and the Social Withdrawal factors correlated with other factors in the Negative Valence Systems domain, rather than the RDoC-assigned grouping of these factors (Cognitive Systems: Attention, and Social Systems: Affiliation & Attachment, respectively). A previous review found that inattention and anxiety are tightly linked (51) but our results provide evidence of the direct link between inattention and negative affect. We also found that the Delay Discounting factor, while considered part of the RDoC Positive Valence System: Reward Valuation subconstruct (52), correlated instead with Cognitive Systems factors, suggesting delay discounting is more related to Cognitive Control/Impulsivity domains than to Reward Valuation (53–55).

**Table 2.**
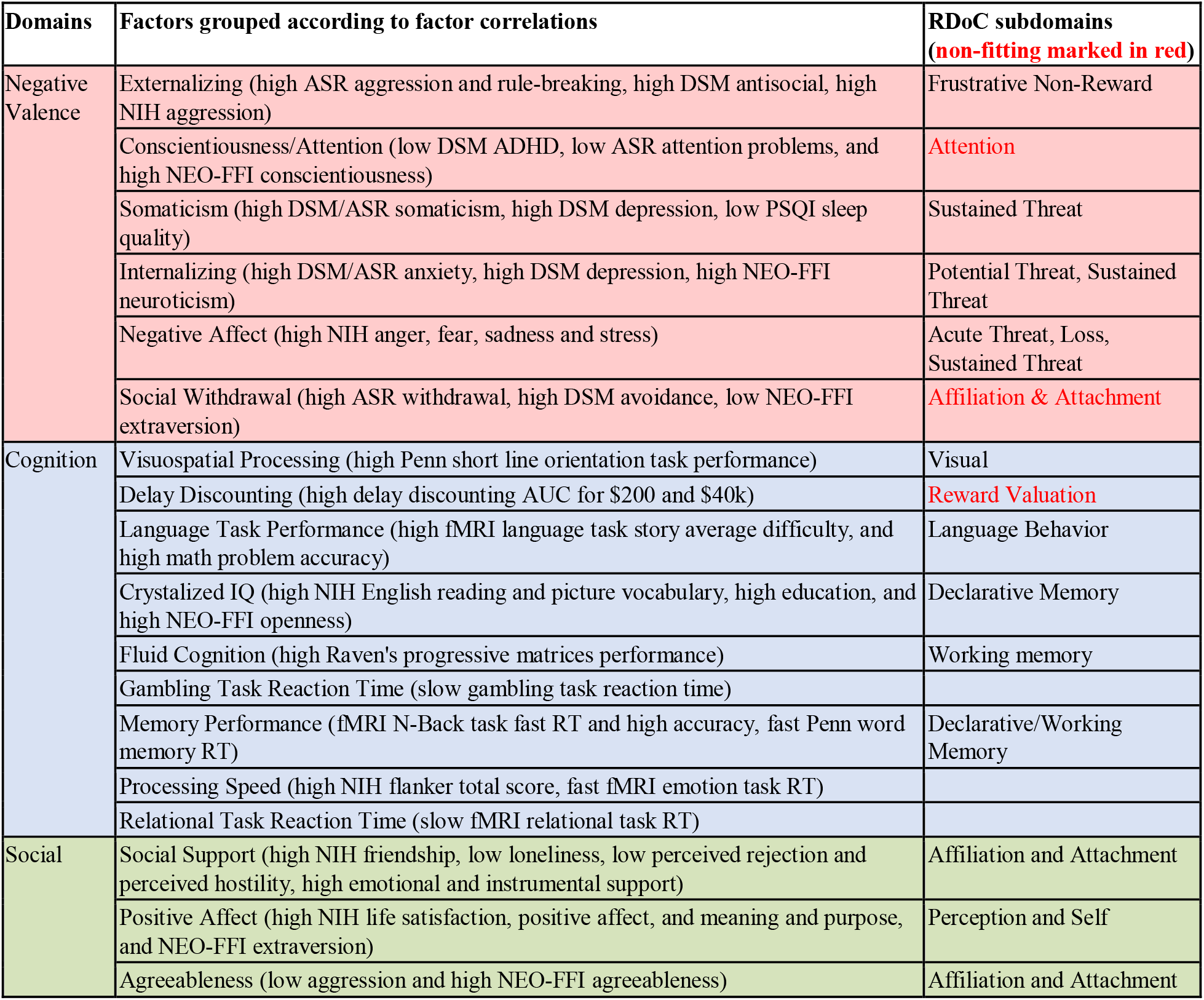
Discovered factors (EFA) using 100 phenotypic measures. Factors are grouped according to correlations between the factors. The right column indicates the RDoC domain each factor most closely approximated. Factors whose correlation structure did not match the RDoC domain assignment for that factor (n = 3) are displayed in red.

Here we summarize several key points from our mapping of causal influences on AUD symptom severity onto the RDoC framework. First, our analysis uncovered strong evidence for the direct causal effect of the Negative Valence domain in AUD. This specifically included a causal influence of the Negative Affect factor on AUD, mediated through Conscientiousness/Attention (which correlated with other Negative Valence Systems factors). While many neurobiological models of addiction agree on the importance of negative affect in AUD (10,20), this is not unanimously agreed upon by experts in the addiction sciences (17). The presented empirical data hence provides important empirical evidence implicating the broader Negative Affect Domain (as defined in RDoC) as an important treatment target in AUD.

Second, our analysis uncovered strong evidence for a mediating/buffering role of the Social Systems domain in AUD. Low Social Support and low Agreeableness were indirect causes of AUD severity and fully mediated the effect of cognition on the negative valence domain, providing strong empirical evidence that addiction models should incorporate measures of social function (56,57). Epidemiological research has established a solid link between social affiliation and drug addiction (58), and increased social affiliation is associated with decreased risk of relapse in drug users who are seeking treatment (59). Despite the considerable evidence research has uncovered for the importance of social affiliation as a protective factor in addiction (60), current neurobiological models of addiction generally fail to consider social factors (56) and their close relationship to cognitive/affective factors.

We generally found weak evidence for the involvement of the Positive Valence Systems domain in this analysis, although this is likely due to a limitation of the dataset employed. Positive Valence Systems subdomains, including reward-based domains that are particularly important in addiction (17) are relatively neglected in the HCP dataset (61). We did find a causal influence of Delay Discounting on AUD severity, but as noted previously the Delay Discounting factor correlated with other Cognitive factors, suggesting that Delay Discounting (as measured in the HCP study) might be related to Cognitive Control/Impulsivity more so than Reward Valuation (53–55).

Finally, our analysis provides strong evidence that prefrontal cortex (PFC) brain networks, and associated cognitive factors, are situated at the top of the causal hierarchy of influences on AUD severity. The role of prefrontal cortex and associated high-level cognitive factors in addiction is often referenced and is a part of major current theories of addiction (20,62), but our results are among the first to empirically demonstrate this hierarchical influence on AUD. Importantly, our causal model indicates that cognitive influences on AUD severity may extend far beyond the traditional consensus that inhibitory control is the most important cognitive influence on addiction (17,62,63), as we also highlighted influences of fluid and Crystallized IQ, working memory, and language on AUD.

### 4.2 Impact of Results on Theorized Externalizing and Internalizing Pathways to AUD

Previous research has shown that externalizing symptomatology predicts AUD in young adult samples (64). Our data-driven causal model revealed that externalizing fully mediated the impact of all other (measured) causes of AUD; that is, AUD is unrelated to other phenotypic or brain network factors when externalizing is controlled for. Note that our externalizing factor consisted of ASR rule-breaking, aggression, and antisocial scales, and NIH toolbox aggression. The ASR rule-breaking scale contains an item that assesses whether a subject “gets drunk,” but this does not appear to have influenced the current analysis. First, this is only one out of 14 items on the rule-breaking scale (and 40 items in total that contributed to the Externalizing factor). Second, all four scales that formed the Externalizing factor independently correlated with AUD symptom severity (all *p* < .001), such that each individual aspect of Externalizing appears to be related to AUD severity. Overall the model hence supports that externalizing symptoms in general mediate the causal influence of other factors on AUD.

Research has also focused on the high coincidence of AUD and internalizing disorders (65–67). Previous causal modeling research found a causal path from internalizing disorder to drinking behavior in AUD (mediated through drinking-to-cope) (68). The current model contains a confounded relationship between Internalizing and Conscientiousness/Attention, indicating an inability of the model to determine the relationship between these two factors, possibly due to underlying constructs (e.g. drinking motives) that were not captured in this data set. Therefore, it remains to be further described by future research if negative affect is a common underlying cause of internalizing and AUD symptoms (65), or if there is an independent causal influence of internalizing psychopathology on AUD symptoms. It is possible that this relationship could be better examined through a longitudinal study, as pathodevelopmental perspectives on AUD have proposed that early stages of addiction are characterized by low levels of internalizing, but later stages of addiction are characterized by increasing levels of internalizing (65).

### 4.3 A Gray Area in the Literature – the Role of Fluid Cognition in AUD

Executive function is central to neurobiological models of addiction (10,20,62,63). Fluid cognition (here, measured through performance on an abbreviated form of Raven’s progressive matrices) (69), or a person’s ability to reason and think abstractly and flexibly, has an intuitive relationship with the concept of executive function. Authors have often considered working memory to be either indicative of executive function (70), or of fluid cognition (71), and executive function and fluid cognition are similarly impacted by brain lesions (72). Our results generate the hypothesis that fluid cognition and Crystalized IQ, including problem-solving and abstract reasoning, lie at the beginning of a causal hierarchy eventually influencing AUD severity. This fits previous empirical evidence demonstrating that AUD is associated with deficits attributed to various fluid cognitive abilities or executive function such as working memory (73,74) and planning and goal maintenance (75), but expands on this by indicating that these factors have a causal influence on AUD symptom severity. Previous research has also demonstrated that high-level cognition predicts initiation of substance use in adolescence (76), lifetime drug use and abuse (77) and addiction treatment outcomes (78). Our model thus adds to the growing body of empirical evidence that proposes a causal role of cognition as a primary resilience factor and potential treatment target in AUD.

### 4.4 Brain Network Influences in AUD

We found a direct brain-phenotype link from frontoparietal (executive) network connectivity to fluid cognition, corroborating previous evidence of this link in healthy populations (79–83). In addiction, frontoparietal network dysfunction has been implicated in impaired inhibitory control (20,84,85) and self-regulation (51,85). Individuals with AUD show decreased recruitment of frontoparietal network during social-emotional processing (86,87), decision-making (88), and cognitive control (89), as well as decreased frontoparietal connectivity during rest (90). Frontoparietal disengagement during social-emotional processing predicts relapse in AUD (91), and decreased frontoparietal activation during inhibitory control predicts later drinking in adolescents (92,93). Our results indicate that causal influences of frontoparietal network connectivity on AUD are mediated through deficits in overall cognitive ability and its downstream effects on the broader Negative Affect domain. A recent systematic review (94) also showed that targeting dlPFC (part of the frontoparietal network) (24) can improve cognitive deficits in addiction, including executive functions. This is particularly relevant, as our data-driven model also indicates that neuromodulation of frontoparietal network could improve executive functioning, with downstream effects on AUD severity.

We also found direct effects of cingulo-opercular and dorsal attention network connectivity onto the frontoparietal network. The Cingulo-opercular network reacts to salient stimuli regardless of positive/negative valence (95,96), while the dorsal attention network supports the external focusing of attention (97) and encodes top-down control and working memory load (98). Individuals with AUD also show decreased cingulo-opercular activation during social-emotional processing (86,87), cognitive control (89,99,100), and decision-making (88). Our results suggest that dysfunctional connectivity in salience and attentional networks can contribute to cognitive dysfunction in AUD, with these effects being mediated through executive network connectivity. The presented causal model hence provides direct evidence for brain-directed treatment approaches targeted at the frontoparietal network, such as cognition-enhancing therapy (85,101,102), pharmacological interventions (cognitive enhancers) (101) or neuromodulation treatment (e.g., by external devices) (94,103) or neurofeedback interventions (51).

A novel result generated from our causal discovery analysis is the role of language network connectivity as a central “hub” in the brain during resting-state. Language network connectivity caused cingulo-opercular network connectivity directly, and indirectly caused dorsal attention network connectivity (mediated through cingulo-opercular connectivity, and posterior multi-modal association network connectivity). Therefore, language network connectivity exerts influences on frontoparietal network connectivity through multiple different pathways, and might have long-range impacts on cognition and eventually on AUD severity. The potential involvement of language network in AUD appears to have been scarcely investigated. This network encompasses vlPFC regions that are close to left dlPFC regions that are often targeted in neuromodulation interventions for addiction (103), and therefore neuromodulation targeted at left DLPFC might also stimulate language networks. Left vlPFC regions are also implicated in cognitive interventions for addiction (85). Future analysis should examine the relationship between brain language networks and cognitive dysfunction in AUD, and the implications for treatment.

### 4.6 Limitations

The analysis method used in the current manuscript is not free of limitations, and other limitations are also imposed by the nature of the dataset we used. Notably, the fact that we did not find any cycles (i.e. variable X causes variable Y, and variable Y causes variable X) in the current data does not mean that they do not exist. The causal discovery algorithm used in this analysis cannot discover recurring cycles in cross-sectional data, but is capable of discovering recurrent cycles when more than one time point is measured and the cycles unfold over time. Future analysis should incorporate longitudinal data to specifically test the possibility that recurrent cycles might contribute to AUD (10). This limitation extends to the brain network subgraph we recovered as well; the causal discovery algorithm we used cannot recover bidirectional relationships in cross-sectional data, so some brain network links that are actually bidirectional processing streams may instead be represented as the predominant causal relationship between two networks. Finally, the CDA algorithm also uses a penalized likelihood score, therefore potentially missing weak causal links present in the data; however, this practice also serves to increase the confidence in the causal relationships the algorithm does find.

An important limitation of the data set is the extremely limited assessment of the Positive Valence domain in the HCP dataset. Current perspectives in addiction emphasize the role of Positive Valence domains (10,17,20,62,63), but the HCP dataset does not contain many measures in this domain. The data did contain a measure of Delay Discounting, which had causal influences on AUD severity, but this factor appeared to be grouped with other cognitive factors and could not be interpreted as a unique measure of Reward Valuation. The HCP dataset also contained a gambling choice fMRI task, but this task did not provide a phenotypic measure of incentive salience processing. Future analysis will need to carefully measure Positive Valence domains, in addition to the domains measured in the HCP dataset, to determine where these domains fit in an overall causal model of AUD. Another limitation of the dataset is that the cross-sectional design employed by the HCP is also unable to assess certain predictions of pathodevelopmental perspectives on addiction, such as the possibility that different causal factors are involved in early and late stages of AUD (65).

### 4.6 Conclusions

This study is the first to conduct a machine learning search for causal influences of AUD symptoms over a wide phenotypic and neurobiological space. We found phenotypic factors related to several RDoC domains, and confirmed hypothesized influences of a Negative Valence (Negative affect > Conscientiousness/Inattention > Externalizing > AUD symptom severity), and Cognitive Systems (Fluid Cognition > Crystalized IQ > Working Memory/Language> Social/Affective/Psychiatric factors > AUD symptom severity) on AUD. The model proposed a hierarchy with causal influence propagating from brain function to cognition (Fluid/Crystalized Cognition, Language & Working Memory) to social (Agreeableness/Social Support) to affective/psychiatric function (Negative Affect, low Conscientiousness/Attention, Externalizing symptoms) and ultimately AUD symptoms. These results underscore a) a strong causal link between prefrontal brain function/cognition and affective/psychiatric factors and b) an important buffer function of social factors (Social Support, Agreeableness). Our data-driven model hence confirmed hypothesized influences of cognitive and affective factors on AUD, while underscoring that traditional addiction models need to be expanded to highlight the importance of social factors, amongst others. Results further demonstrated that it is possible to reduce a broad phenotypic space (100 measures) to a limited set of causal factors of AUD, which can inform future research. We argue that the presented causal model of AUD provides evidence for exploring two different kinds of treatment approaches, specifically for investigating a) “top-down” interventions aimed at enhancing high-level cognition, including brain-directed interventions targeting the executive network and b) “integrative” interventions that take the interplay between brain/cognitive, affective/psychiatric factors and social factors into account. We note that we did not investigate the individual heterogeneity of the causal factors involved in this model, but only provided a static causal model of an “average” individual with AUD symptoms, as a first step. We believe that this initial step of describing a comprehensive, integrated, multimodal but also reduced model (in a principled data-driven way) is crucial. We see the provided causal model as a working model, which can be further expanded (e.g. by the RDoC Positive Valence factors), explored with regards to individual heterogeneity and used in predictive modelling studies on alcohol use trajectories in active users, as well as in individuals with AUD in treatment.

## 5 Acknowledgments

ER is supported by National Institutes of Mental Health (NIMH) grant T32-MH115866 to David Redish. EK received support for this work from the National Center for Advancing Translational Sciences of the National Institutes of Health Award Number UL1TR000114. The content is solely the responsibility of the authors and does not necessarily represent the official views of the National Institutes of Health or the National Institutes of Mental Health. The authors thank Matthew Kushner for valuable input and editorial assistance.

