## Supplement for "An Integrated Multimodal Model of Alcohol Use Disorder Generated by Data-Driven Causal Discovery Analysis"

**Supplementary Tables:**

*Table S1. List of included variables and excluded variables.*

| **Supplementary Table S1. Variables included in factor analysis of the HCP dataset.** | |
| --- | --- |
| **Self-Report Psychiatric Symptoms** |  |
| ASR_Witd_T | Withdrawal |
| ASR_Anxd_T | Anxiety (ASR) |
| ASR_Soma_T | Somaticism (ASR) |
| ASR_Thot_T | Disruptive Thoughts |
| ASR_Attn_T | Attention Deficits |
| ASR_Aggr_T | Aggression |
| ASR_Rule_T | Rule Breaking |
| ASR_Intr_T | Intrusive thoughts |
| DSM_Depr_T | Depression |
| DSM_Anxi_T | Anxiety (DSM) |
| DSM_Somp_T | Somaticism (DSM) |
| DSM_Avoid_T | Avoidance |
| DSM_Adh_T | ADHD Symptoms |
| DSM_Antis_T | Antisocial |
| SSAGA_ChildhoodConduct | Childhood Conduct Disorder |
| SSAGA_Depressive_Sx | Depressive Symptoms |
| **NIH Negative Affect Toolbox** |  |
| AngAffect_Unadj | Angry affect |
| AngHostil_Unadj | Anger (hostility) |
| AngAggr_Unadj | Anger (aggression) |
| FearAffect_Unadj | Fearful affect |
| FearSomat_Unadj | Fear (somaticism) |
| Sadness_Unadj | Sadness |
| **NIH Social Relationships Toolbox** |  |
| Friendship_Unadj | Friendship |
| Loneliness_Unadj | Loneliness |
| PercHostil_Unadj | Perceived Hostility |
| PercReject_Unadj | Perceived Rejection |
| EmotSupp_Unadj | Emotional Support |
| InstruSupp_Unadj | Instrumental Support |
| PercStress_Unadj | Perceived Stress |
| **Penn Emotion Recognition Test** |  |
| ER40_CRT | Reaction Time (correct) |
| ER40ANG | Anger recognition |
| ER40FEAR | Fearful recognition |
| ER40HAP | Happy recognition |
| ER40NOE | No emotion recognition |
| ER40SAD | Sadness recognition |
| **Demographics** |  |
| SSAGA_Income | Income bracket |
| SSAGA_Educ | Educational level attained |
| **In-Scanner Behavior (Gambling Task)** |  |
| Gambling_Task_Reward_Perc_Larger | Gambling task reward % guessed larger |
| Gambling_Task_Reward_Median_RT_Larger | Gambling task average RT (reward-larger) |
| Gambling_Task_Reward_Median_RT_Smaller | Gambling task average RT (reward-smaller) |
| Gambling_Task_Punish_Perc_Larger | Gambling task punish % guessed larger |
| Gambling_Task_Punish_Median_RT_Larger | Gambling task average RT (punish-larger) |
| Gambling_Task_Punish_Median_RT_Smaller | Gambling task average RT (punish-smaller) |
| **In-Scanner Behavior (Relational Task)** |  |
| Relational_Task_Match_Acc | Relational Task Accuracy (Matching) |
| Relational_Task_Match_Median_RT | Relational Task Median RT (Matching) |
| Relational_Task_Rel_Acc | Relational Task Accuracy (Relational) |
| Relational_Task_Rel_Median_RT | Relational Task Median RT (Relational) |
| **In-Scanner Behavior (Emotion task)** |  |
| Emotion_Task_Face_Acc | Emotion task accuracy (face) |
| Emotion_Task_Face_Median_RT | Emotion task median RT (face) |
| Emotion_Task_Shape_Acc | Emotion task accuracy (shape) |
| Emotion_Task_Shape_Median_RT | Emotion task median RT (shape) |
| **Fluid Intelligence** |  |
| PMAT24_A_CR | Raven's Progressive Matrices - # Correct |
| PMAT24_A_SI | Raven's Progressive Matrices - # Skipped |
| PMAT24_A_RTCR | Raven's Progressive Matrices - Reaction Times (Correct Only) |
| **Delay Discounting** |  |
| DDisc_AUC_200 | Delay Discounting AU-ROC ($200) |
| DDisc_AUC_40K | Delay Discounting AU-ROC ($40k) |
| **NIH Cognition Toolbox** |  |
| PicSeq_AgeAdj | Picture Sequence Memory (episodic memory) |
| CardSort_AgeAdj | Dimensional Change Card Sorting (cognitive flexibility) |
| Flanker_AgeAdj | Flanker Task (attention/inhibitory control) |
| ReadEng_AgeAdj | English reading ability |
| PicVocab_AgeAdj | Picture Vocabulary |
| ProcSpeed_AgeAdj | Pattern Completion Processing Speed |
| ListSort_AgeAdj | List Sorting (Working Memory) |
| **NIH Psychological Well-Being Toolbox** |  |
| LifeSatisf_Unadj | General Life Satisfaction |
| MeanPurp_Unadj | Meaning and Purpose in Life |
| PosAffect_Unadj | Positive Affect |
| **Visuospatial Processing (Variable Short Continuous Penn Line Orientation Test)** |  |
| VSPLOT_TC | Correct Responses |
| VSPLOT_CRTE | Reaction time divided by expected clicks |
| VSPLOT_OFF | Total positions off for each trial |
| **Attention (Short Continuous Penn Test)** |  |
| SCPT_SEN | SCPT Sensitivity |
| SCPT_SPEC | SCPT Specificity |
| **Penn Word Memory Test** |  |
| IWRD_TOT | Total # of correct responses |
| IWRD_RTC | Reaction time for correct responses |
| **Sensory** |  |
| Noise_Comp | Noise sensitivity |
| Odor_AgeAdj | Odor sensitivity |
| PainInterf_Tscore | Pain intensity and interference |
| Taste_AgeAdj | Taste sensitivity |
| Mars_Final | Visual contrast sensitivity |
| **In-Scanner Behavior (Working Memory [N-back])** |  |
| WM_Task_2bk_Acc | Working memory task accuracy (2-back) |
| WM_Task_2bk_Median_RT | Working memory task RT (2-back) |
| WM_Task_0bk_Acc | Working memory task accuracy (0-back) |
| WM_Task_0bk_Median_RT | Working memory task RT (0-back) |
| **In-Scanner Behavior (Social Task)** |  |
| Social_Task_Perc_Random | Social task % guessed random |
| Social_Task_Perc_TOM | Social task % guessed TOM |
| Social_Task_Median_RT_Random | Social task Median RT on Random |
| Social_Task_Median_RT_TOM | Social Task Median RT on TOM |
| **In-Scanner Behavior (Language task)** |  |
| Language_Task_Story_Acc | Language task accuracy (story) |
| Language_Task_Story_Median_RT | Language task median RT (story) |
| Language_Task_Story_Avg_Difficulty_Level | Language task average difficulty (story) |
| Language_Task_Math_Acc | Language task accuracy (math) |
| Language_Task_Math_Median_RT | Language task median RT (math) |
| Language_Task_Math_Avg_Difficulty_Level | Language task average difficulty (math) |

*Table S2: Descriptive statistics of the included variables.*

| **Variable** | **mean** | **sd** |
| --- | --- | --- |
| ASR_Witd_T | 53.5 | 5.8 |
| ASR_Anxd_T | 53.9 | 6.3 |
| ASR_Soma_T | 53.9 | 5.9 |
| ASR_Thot_T | 53.7 | 5.7 |
| ASR_Attn_T | 55 | 5.7 |
| ASR_Aggr_T | 52.5 | 4.1 |
| ASR_Rule_T | 53.8 | 5.2 |
| ASR_Intr_T | 53.8 | 5.4 |
| DSM_Depr_T | 53.9 | 5.8 |
| DSM_Anxi_T | 53.3 | 5.3 |
| DSM_Somp_T | 54 | 5.7 |
| DSM_Avoid_T | 54.5 | 6.1 |
| DSM_Adh_T | 54.9 | 5.9 |
| DSM_Antis_T | 52.9 | 4.7 |
| SSAGA_ChildhoodConduct | 0.5 | 0.8 |
| SSAGA_Depressive_Sx | 1.3 | 2.5 |
| MMSE_Score | 29 | 1 |
| PSQI_Score | 4.7 | 2.7 |
| PMAT24_A_CR | 17 | 4.7 |
| PMAT24_A_SI | 2.9 | 3.8 |
| PMAT24_A_RTCR | 15943.5 | 9241.8 |
| DDisc_AUC_200 | 0.3 | 0.2 |
| DDisc_AUC_40K | 0.5 | 0.3 |
| PicSeq_AgeAdj | 105.4 | 16.4 |
| CardSort_AgeAdj | 102.6 | 9.7 |
| Flanker_AgeAdj | 102.1 | 9.9 |
| ReadEng_AgeAdj | 107.2 | 14.6 |
| PicVocab_AgeAdj | 109.4 | 14.8 |
| ProcSpeed_AgeAdj | 104.3 | 19.8 |
| ListSort_AgeAdj | 103.7 | 13.2 |
| AngAffect_Unadj | 47.5 | 8.2 |
| AngHostil_Unadj | 50.5 | 8.6 |
| AngAggr_Unadj | 51.9 | 8.8 |
| FearAffect_Unadj | 50 | 7.9 |
| FearSomat_Unadj | 51.7 | 8.2 |
| Sadness_Unadj | 46 | 7.8 |
| LifeSatisf_Unadj | 54.7 | 9.3 |
| MeanPurp_Unadj | 52 | 8.8 |
| PosAffect_Unadj | 50.3 | 7.8 |
| Friendship_Unadj | 50.5 | 9 |
| Loneliness_Unadj | 50.9 | 8.6 |
| PercHostil_Unadj | 48.6 | 8.6 |
| PercReject_Unadj | 48.4 | 8.7 |
| EmotSupp_Unadj | 51.5 | 9.5 |
| InstruSupp_Unadj | 48.2 | 8.9 |
| PercStress_Unadj | 48.1 | 9.1 |
| SelfEff_Unadj | 51.1 | 8.3 |
| VSPLOT_TC | 15 | 4.4 |
| VSPLOT_CRTE | 1156.2 | 356.1 |
| VSPLOT_OFF | 23.7 | 14.2 |
| SCPT_SPEC | 1 | 0 |
| SCPT_SEN | 1 | 0.1 |
| IWRD_TOT | 35.6 | 2.9 |
| IWRD_RTC | 1558 | 296.8 |
| ER40_CRT | 1829.9 | 320.9 |
| ER40ANG | 6.8 | 1 |
| ER40FEAR | 6.9 | 1.2 |
| ER40NOE | 7.1 | 1.2 |
| ER40SAD | 6.8 | 1.1 |
| ER40HAP | 8 | 0.2 |
| NEOFAC_A | 33.7 | 5.8 |
| NEOFAC_O | 28.4 | 6.2 |
| NEOFAC_C | 34.5 | 5.9 |
| NEOFAC_N | 16.5 | 7.4 |
| NEOFAC_E | 30.8 | 6 |
| PainInterf_Tscore | 45.5 | 7.5 |
| Noise_Comp | 4.4 | 1.5 |
| Odor_AgeAdj | 98.1 | 11.2 |
| Taste_AgeAdj | 93.7 | 14.6 |
| Mars_Final | 1.8 | 0.6 |
| Emotion_Task_Face_Acc | 98.4 | 3.6 |
| Emotion_Task_Face_Median_RT | 786.4 | 132 |
| Emotion_Task_Shape_Acc | 96.6 | 4.4 |
| Emotion_Task_Shape_Median_RT | 766 | 114 |
| Language_Task_Story_Acc | 94.3 | 8.5 |
| Language_Task_Story_Median_RT | 3303.9 | 328.4 |
| Language_Task_Story_Avg_Difficulty_Level | 10.4 | 1.5 |
| Language_Task_Math_Acc | 83.8 | 9.8 |
| Language_Task_Math_Median_RT | 3799.7 | 310 |
| Language_Task_Math_Avg_Difficulty_Level | 2.6 | 0.6 |
| Relational_Task_Match_Acc | 87 | 11.9 |
| Relational_Task_Match_Median_RT | 1446.8 | 256.8 |
| Relational_Task_Rel_Acc | 65.7 | 17.8 |
| Relational_Task_Rel_Median_RT | 1994 | 425.9 |
| Gambling_Task_Reward_Perc_Larger | 50.9 | 13 |
| Gambling_Task_Reward_Median_RT_Larger | 416.2 | 116.8 |
| Gambling_Task_Reward_Median_RT_Smaller | 422 | 118.9 |
| Gambling_Task_Punish_Perc_Larger | 51.2 | 10.8 |
| Gambling_Task_Punish_Median_RT_Larger | 398.4 | 119.2 |
| Gambling_Task_Punish_Median_RT_Smaller | 404.4 | 118.3 |
| WM_Task_2bk_Acc | 83.8 | 10.5 |
| WM_Task_2bk_Median_RT | 963.2 | 143.3 |
| WM_Task_0bk_Acc | 90.4 | 10.3 |
| WM_Task_0bk_Median_RT | 762.7 | 129.3 |
| Social_Task_Perc_Random | 44.3 | 9.9 |
| Social_Task_Perc_TOM | 49 | 7.7 |
| Social_Task_Median_RT_Random | 1059.8 | 366.2 |
| Social_Task_Median_RT_TOM | 1014.4 | 290.2 |
| SSAGA_Income | 5.1 | 2.1 |
| SSAGA_Educ | 15 | 1.8 |

*Table S3: Parallel Analysis to determine number of factors for EFA.*

| Factor Number | Eigenvalue | Random | Resample |
| --- | --- | --- | --- |
| **1** | **13.35512352** | **0.74452256** | **0.745954763** |
| **2** | **7.282829607** | **0.685166489** | **0.685141491** |
| **3** | **4.016473844** | **0.65164259** | **0.651598264** |
| **4** | **2.620050736** | **0.623015644** | **0.622993935** |
| **5** | **1.80891203** | **0.597485868** | **0.597311753** |
| **6** | **1.614116725** | **0.573749405** | **0.573747976** |
| **7** | **1.20109115** | **0.551629253** | **0.551789052** |
| **8** | **1.118315762** | **0.530825242** | **0.530813878** |
| **9** | **1.017203768** | **0.51077243** | **0.510796257** |
| **10** | **0.796850334** | **0.491859768** | **0.491770522** |
| **11** | **0.735685554** | **0.473513162** | **0.473437437** |
| **12** | **0.689950612** | **0.455839456** | **0.455628477** |
| **13** | **0.595127371** | **0.438724077** | **0.43847865** |
| **14** | **0.5808096** | **0.422019326** | **0.421887444** |
| **15** | **0.493719289** | **0.405737228** | **0.405638205** |
| **16** | **0.468799252** | **0.389937853** | **0.389931539** |
| **17** | **0.413306447** | **0.374503656** | **0.374684555** |
| **18** | **0.400562751** | **0.359629076** | **0.35967957** |
| *19* | *0.331990827* | *0.344669984* | *0.344905954* |
| *20* | *0.297733166* | *0.33041483* | *0.330339106* |
| *21* | *0.277038289* | *0.316310885* | *0.316209627* |
| *22* | *0.238786103* | *0.302283719* | *0.302288296* |
| *23* | *0.214286433* | *0.288669245* | *0.288614506* |
| *24* | *0.204540862* | *0.27519919* | *0.275267838* |
| *25* | *0.196761536* | *0.261832044* | *0.261950458* |
| *26* | *0.141291263* | *0.248801588* | *0.248922675* |
| *27* | *0.121906185* | *0.236033707* | *0.236104985* |
| *28* | *0.109101529* | *0.223366786* | *0.223530283* |
| *29* | *0.103150058* | *0.210887547* | *0.210960733* |
| *30* | *0.079015654* | *0.198574338* | *0.198674535* |
| *31* | *0.045756892* | *0.186434393* | *0.186502655* |
| *32* | *0.021814247* | *0.174373956* | *0.174396029* |
| *33* | *-0.009245324* | *0.162485744* | *0.162516646* |
| *34* | *-0.016100507* | *0.150755491* | *0.15074531* |
| *35* | *-0.040693603* | *0.139180742* | *0.1391719* |
| *36* | *-0.045133732* | *0.127717866* | *0.127673068* |
| *37* | *-0.06195933* | *0.11636482* | *0.11631437* |
| *38* | *-0.084790592* | *0.105110434* | *0.105153228* |
| *39* | *-0.101181919* | *0.093995307* | *0.093988958* |
| *40* | *-0.114578626* | *0.083035864* | *0.082975901* |
| *41* | *-0.132123879* | *0.072094566* | *0.072057412* |
| *42* | *-0.138903842* | *0.061171723* | *0.061210195* |
| *43* | *-0.145098865* | *0.050503478* | *0.050386157* |
| *44* | *-0.17124058* | *0.039848126* | *0.039737408* |
| *45* | *-0.176370681* | *0.029176857* | *0.029206826* |
| *46* | *-0.186645179* | *0.018689989* | *0.018712451* |
| *47* | *-0.193683698* | *0.008248094* | *0.00828573* |
| *48* | *-0.204564542* | *-0.002062978* | *-0.002016905* |
| *49* | *-0.214627909* | *-0.012161367* | *-0.012187036* |
| *50* | *-0.226758407* | *-0.022415891* | *-0.022370874* |
| *51* | *-0.240490081* | *-0.03255589* | *-0.032522555* |
| *52* | *-0.250712622* | *-0.042511608* | *-0.042519157* |
| *53* | *-0.262379255* | *-0.052422081* | *-0.052458683* |
| *54* | *-0.273476241* | *-0.062486479* | *-0.062355672* |
| *55* | *-0.301753867* | *-0.072401112* | *-0.072298804* |
| *56* | *-0.311253717* | *-0.08216437* | *-0.08207893* |
| *57* | *-0.323096077* | *-0.09184905* | *-0.091853034* |
| *58* | *-0.336840527* | *-0.10151253* | *-0.101467694* |
| *59* | *-0.338225145* | *-0.111175097* | *-0.111182147* |
| *60* | *-0.34825169* | *-0.120750619* | *-0.12075821* |
| *61* | *-0.350060561* | *-0.130161066* | *-0.13036507* |
| *62* | *-0.358412062* | *-0.139798829* | *-0.139765705* |
| *63* | *-0.359334315* | *-0.149258278* | *-0.149183386* |
| *64* | *-0.373098643* | *-0.158656591* | *-0.158565465* |
| *65* | *-0.382064832* | *-0.168061647* | *-0.167962169* |
| *66* | *-0.395292719* | *-0.177395214* | *-0.177290921* |
| *67* | *-0.400524846* | *-0.186685575* | *-0.186585863* |
| *68* | *-0.416512849* | *-0.195981441* | *-0.195954175* |
| *69* | *-0.426638043* | *-0.20526288* | *-0.205209746* |
| *70* | *-0.42968121* | *-0.214521554* | *-0.214435183* |
| *71* | *-0.441868423* | *-0.223729865* | *-0.223724938* |
| *72* | *-0.453567962* | *-0.232893156* | *-0.232947565* |
| *73* | *-0.458667937* | *-0.242104665* | *-0.242100626* |
| *74* | *-0.468629225* | *-0.251306326* | *-0.25131472* |
| *75* | *-0.478505647* | *-0.26054965* | *-0.260488023* |
| *76* | *-0.496056734* | *-0.269687561* | *-0.269655322* |
| *77* | *-0.499683481* | *-0.278887487* | *-0.278863578* |
| *78* | *-0.518261684* | *-0.28802116* | *-0.288091923* |
| *79* | *-0.523551078* | *-0.297151421* | *-0.297228976* |
| *80* | *-0.533592436* | *-0.306404897* | *-0.306537169* |
| *81* | *-0.552530295* | *-0.315630902* | *-0.315780231* |
| *82* | *-0.55463231* | *-0.324892537* | *-0.325061507* |
| *83* | *-0.566534666* | *-0.334233856* | *-0.334217955* |
| *84* | *-0.576908864* | *-0.343631984* | *-0.343656751* |
| *85* | *-0.586861774* | *-0.353111924* | *-0.353071311* |
| *86* | *-0.592600927* | *-0.362580356* | *-0.362554446* |
| *87* | *-0.641860484* | *-0.372111612* | *-0.371982711* |
| *88* | *-0.666151933* | *-0.38173192* | *-0.381621584* |
| *89* | *-0.703993759* | *-0.391462812* | *-0.391413423* |
| *90* | *-0.710154898* | *-0.401303495* | *-0.401281033* |
| *91* | *-0.744792303* | *-0.4113873* | *-0.411331228* |
| *92* | *-0.749874482* | *-0.421568213* | *-0.421535219* |
| *93* | *-0.761787715* | *-0.432066018* | *-0.432075655* |
| *94* | *-0.773111592* | *-0.442790436* | *-0.442938688* |
| *95* | *-0.786254514* | *-0.45385102* | *-0.454152578* |
| *96* | *-0.803679447* | *-0.465534192* | *-0.465715286* |
| *97* | *-0.819855771* | *-0.477879841* | *-0.47797266* |
| *98* | *-0.838758759* | *-0.491335741* | *-0.491337841* |
| *99* | *-0.846681015* | *-0.506401219* | *-0.50639727* |
| *100* | *-0.945766866* | *-0.52578785* | *-0.525694368* |

*Figure S1. Permutation test for eigenvalue significance (parallel analysis).*

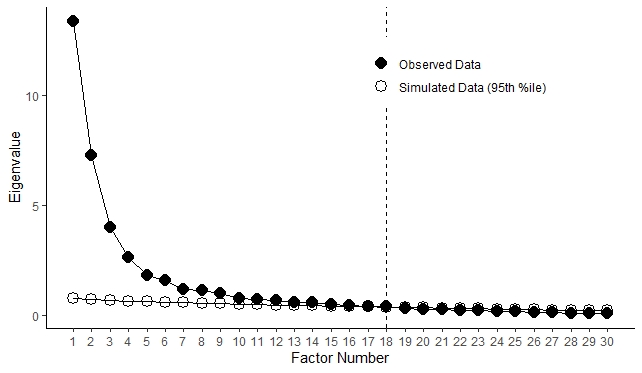

*Table S4. Factor loadings of the included variables.*

This table is too large to be properly displayed in a word document; this table is instead provided as a spreadsheet for download.

*Table S5. Percent variance explained per factor, eigenvalues, and cumulative variance.*

| **Factor** | **Proportion Variance** | **Cumulative Variance** | **Eigenvalue** |
| --- | --- | --- | --- |
| Somaticism | 0.03 | 0.03 | 13.36 |
| Fluid Cognition | 0.03 | 0.06 | 7.28 |
| Internalizing | 0.03 | 0.09 | 4.02 |
| Gamble RT | 0.03 | 0.12 | 2.62 |
| Inattention | 0.03 | 0.16 | 1.81 |
| Visuospatial | 0.02 | 0.19 | 1.61 |
| Social Support | 0.03 | 0.22 | 1.2 |
| Processing Speed | 0.02 | 0.24 | 1.12 |
| Externalizing | 0.03 | 0.27 | 1.02 |
| Avoidance | 0.02 | 0.29 | 0.8 |
| Language Task | 0.02 | 0.32 | 0.74 |
| Relational Task | 0.02 | 0.35 | 0.69 |
| Delay Discounting | 0.02 | 0.37 | 0.59 |
| N-Back Task | 0.03 | 0.39 | 0.58 |
| Negative Affect | 0.03 | 0.41 | 0.49 |
| CrystalizedIQ | 0.03 | 0.43 | 0.47 |
| Positive Affect | 0.02 | 0.44 | 0.41 |
| Agreeableness | 0.01 | 0.46 | 0.4 |

*Table S6. Factor correlation structure.*

This table is too large to be properly displayed in a word document; this table is instead provided as a spreadsheet for download.

*Table S7. Resampling stability of graph edges.*

| **Var1** | **Var2** | **Stability** |
| --- | --- | --- |
| PosteriorMM | Visual1 | 0.6525 |
| Visual1 | VentralMM | 0.496 |
| Language | VentralMM | 0.831 |
| Visual1 | Auditory | 0.666 |
| Auditory | Somatomotor | 0.785 |
| Language | PosteriorMM | 0.658 |
| VentralMM | Orbitoaffective | 0.9585 |
| CinguloOpercular | Orbitoaffective | 0.9405 |
| Somatomotor | Visual2 | 0.943 |
| DorsalAttention | Visual2 | 0.1605 |
| Language | DefaultMode | 0.858 |
| Frontoparietal | DefaultMode | 0.9325 |
| Language | CinguloOpercular | 0.0555 |
| PosteriorMM | DorsalAttention | 0.691 |
| CinguloOpercular | DorsalAttention | 0.242 |
| CinguloOpercular | Frontoparietal | 0.58 |
| DorsalAttention | Frontoparietal | 0.789 |
| Frontoparietal | FluidCognition | 0.4045 |
| FluidCognition | Visuospatial | 0.638 |
| FluidCognition | CrystallizedIQ | 0.6355 |
| Visuospatial | ProcessingSpeed | 0.723 |
| WorkingMemory | ProcessingSpeed | 0.902 |
| Visuospatial | LanguageTask | 0.6445 |
| CrystallizedIQ | LanguageTask | 0.6145 |
| CrystallizedIQ | DelayDiscounting | 0.9255 |
| ProcessingSpeed | RelationalTask | 0.902 |
| LanguageTask | WorkingMemory | 0.808 |
| LanguageTask | Agreeableness | 0.451 |
| DelayDiscounting | Agreeableness | 0.2485 |
| RelationalTask | GambleRT | 0.2475 |
| WorkingMemory | GambleRT | 0.9025 |
| WorkingMemory | SocialSupport | 0.3335 |
| Agreeableness | SocialSupport | 0.295 |
| Agreeableness | Externalizing | 0.48 |
| Conscientiousness | Externalizing | 0.4685 |
| SocialSupport | Avoidance | 0.9465 |
| Internalizing | Avoidance | 0.8395 |
| SocialSupport | NegativeAffect | 0.453 |
| SocialSupport | PositiveAffect | 0.9355 |
| Externalizing | PositiveAffect | * |
| NegativeAffect | Internalizing | 0.386 |
| NegativeAffect | Conscientiousness | 0.209 |
| Internalizing | Somaticism | 0.4065 |
| Conscientiousness | Somaticism | 0.431 |
| Externalizing | AUD | 0.8195 |
